# Deep learning methods to forecasting human embryo development in time-lapse videos

**DOI:** 10.1101/2024.03.04.583269

**Authors:** Akriti Sharma, Alexandru Dorobantiu, Saquib Ali, Mario Iliceto, Mette H. Stensen, Erwan Delbarre, Michael A. Riegler, Hugo L. Hammer

## Abstract

**Background:** In assisted reproductive technology, evaluating the quality of the embryo is crucial when selecting the most viable embryo for transferring to a woman. Assessment also plays an important role in determining the optimal transfer time, either in the cleavage stage or in the blastocyst stage. Several AI-based tools exist to automate the assessment process. However, none of the existing tools predicts upcoming video frames to assist embryologists in the early assessment of embryos. In this paper, we propose an AI system to forecast the dynamics of embryo morphology over a time period in the future.

**Methods:** The AI system is designed to analyze embryo development in the past two hours and predict the morphological changes of the embryo for the next two hours. It utilizes a predictive model incorporating Convolutional LSTM layers, to predict the future video frame by analyzing prior morphological changes within the embryo’s video sequence. The system uses the predictions recursively and forecasts up to 23 hours of embryo development.

**Results:** The results demonstrated that the AI system could accurately forecast embryo development at the cleavage stage on day 2 and the blastocyst stage on day 4. The system provided valuable information on the cell division processes on day 2 and the start of the blastocyst stage on day 4. The system focused on specific developmental features effective across both the categories of embryos. The embryos that were transferred to the female, and the embryos that were discarded. However, in the ‘transfer’ category, the forecast had a clearer cell membrane and less distortion as compared to the ‘avoid’ category.

**Conclusion:** This study assists in the embryo evaluation process by providing early insights into the quality of the embryo for both the transfer and avoid categories of videos. The embryologists recognize the ability of the forecast to depict the morphological changes of the embryo. Additionally, enhancement in image quality has the potential to make this approach relevant in clinical settings.

**Author summary:** The emergence of assisted reproductive technology has significantly improved infertility treatments. It involves fertilization of an egg outside the body, and the resultant embryos are developed in time-lapse incubators. The embryologists manually evaluate embryos using time-lapse videos and rank each embryo on the basis of several criteria including the dynamics of embryo cell stages, such as the start of the blastocyst stage. Traditional manual analysis is subjective and time-consuming, and AI tools are introduced to automate and enhance embryo selection efficiency. However, current AI tools do not generate video frames that forecast changes in embryo morphology. This study fills this gap by introducing an AI system that forecasts upcoming frames of a time-lapse video. In this approach, several hours were predicted ahead of the last video frame. The system was evaluated on crucial days of embryo evaluation. Our approach was effective in both good quality (transfer) and poor quality (avoid) video categories, and the forecast revealed crucial insights about embryo cell division and the start of the blastocyst stage. Despite some image quality issues, the proposed AI system demonstrated the potential for early and accurate assessment of embryo quality.

## Introduction

The advancement of assisted reproductive technology (ART), including procedures like *in-vitro* fertilization (IVF) and intracytoplasmic sperm injection (ICSI), has provided new possibilities for treating infertility and enhancing the likelihood of conceiving a child with medical treatment [1, 2]. In ART, eggs are retrieved from a woman, fertilized outside the body and the embryos are cultured in incubators. An ‘embryo’ represents the initial stage of human development [3] and is characterized by a sequence of cell divisions referred to as embryo cell stages. Fig 1 shows the development of an embryo spanning across different embryo cell stages. Each division occurs over a time span of 12 to 24 hours, and this progression is measured in hours post insemination (hpi). Initially, the embryo undergoes its first division, resulting in two cells, which then further divide into more cells. As the embryo develops, the cells gradually start to compact together, forming a compact mass known as a morula. Later, these cells begin to differentiate and evolve into a more specific structure called a blastocyst, characterized by distinctive features such as inner cell mass and trophectoderm.

**Fig 1.**
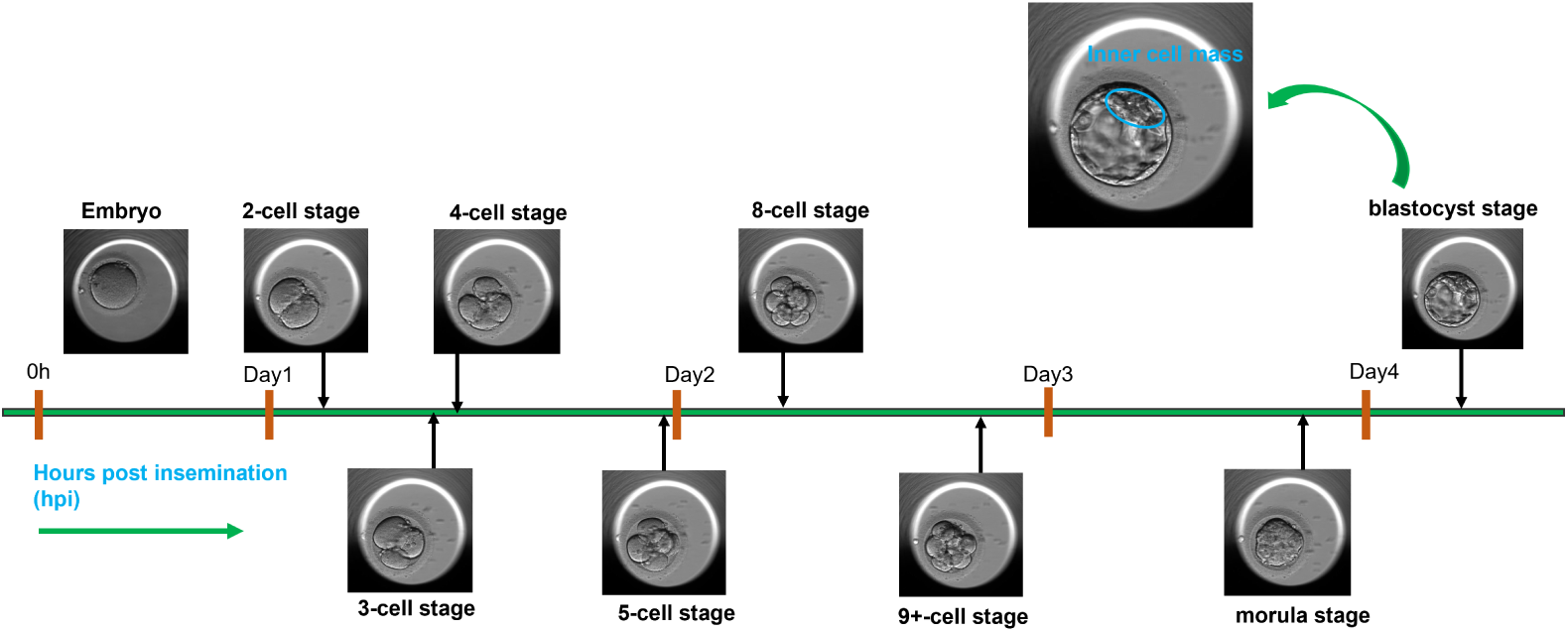
Time-lapse images of embryo development. After fertilization, embryo development spans across different cell stages, morula and blastocyst. The development is grouped into days. The blastocyst stage is magnified to depict the inner cell mass.

Embryos are typically cultured in an incubator until the fifth day of embryo development. Then, an embryo is either transferred to the uterus for implantation, cryopreserved for subsequent transfer or discarded (avoid). This is process is referred to as ‘embryo evaluation’ and is conducted by embryologists after examining time-lapse videos capturing the progression of embryo development during the incubation period [4]. The embryologists base their assessment on several factors including the dynamics of change in embryo morphology [5, 6], the time interval between adjacent cell stages [7, 8] and the start of blastocyst stage [9, 10]. These parameters indicate the implantation potential of an embryo.

Time-lapse incubators facilitate the assessment process by providing embryologists with nearly continuous monitoring of embryo development through frequent image acquisition [11]. These images can be collated into a time-lapse video depicting the entire development process. Traditionally, embryologists manually analyze these videos, a method that is often time-intensive and subjective [12]. To enhance efficiency, various artificial intelligence (AI) algorithms and tools are used within fertility clinics [13].

These AI systems assist embryologists by automating various tasks such as the annotation of cell stages [14, 15], scoring and grading morphology stages [16], embryo selection [4], and the prediction of implantation potential and live-birth outcomes [17–19]. From a broader perspective, these AI systems can be categorized into two types: 1) AI systems that analyze parts or entire time-lapse videos to identify morphological patterns linked to specific outcomes, such as pregnancy, 2) AI systems that analyze the current progression of morphology for predicting the future of embryo development.

In this study, we focus on AI systems that fall into the second category. These AI systems are efficient at predicting the relevant biomarkers, which are crucial for assessing the quality of embryos. A recently introduced AI-based software system, referred to as Cultivating Human Life through Optimal Embryos (*Chloe*) has claimed to digitalize the workflow for embryo selection. It utilizes a combination of various AI models, including convolutional networks (CNNs) for analyzing embryo morphology and patient data, dynamic programming applied to CNNs, temporal action segmentation, and Gardner scoring per video frame, all employed over a long short-term memory (LSTM) network. The models collectively provide early predictions for cell stage detection, expected start of cell stages (in hours post-insemination) as well as predicting and grading the blastocyst stage and the implantation score. These predictions are automatically updated as embryo development progresses, effectively addressing any time discrepancies that might have existed. Also, if clinics are interested in utilizing the tool, they must acquire software licenses, making it cost-intensive.

These systems, however, currently do not generate upcoming frames in a time-lapse video, visually representing developmental changes over subsequent hours. Our approach forecasts embryo development in the following 12 to 23 hours, which allows for an earlier transfer of the embryo to the female. This allows for diminished epigenetic risks and for a reduction in the embryologist’s work load. Specific studies have linked epigenetic risks with prolonged period of embryo incubation [20, 21] and an early assessment can reduce the impact of the modifications. Hence, in this study, we propose an AI model that predicts upcoming frames for time-lapse videos, covering developmental cell stages of the embryo from 31 to 43 hpi (day 2 of embryo development) and from 90 to 113 hpi (day 4 of embryo development). We focused on the training and evaluation of our AI model on day 2 and day 4 embryo development since existing research highlights the significance of the cleavage stage transfer (day 2 to 3) and the blastocyst stage transfer (day 4 to 6) [3, 22]. By concentrating on these specific time interval, we aligned our proposed system with embryo development stages that are relevant for transfer decisions in clinical settings. Furthermore, previous studies on similar AI models observed high quality results around day 2 and day 4 [23]. Consequently, we chose embryo development activities on these particular days as the starting point for our study.

Our suggested AI system employs a forecasting strategy operating on an input video sequence consisting of seven frames to forecast the subsequent seven frames of the sequence depicting potential changes in embryo morphology. So, the system utilized the current morphology dynamics of two hours and forecast two hours in the future. Upon forecasting the last seventh frame, the input sequence shifts by one, incorporating a new frame, and continues to forecast the subsequent embryo’s development. This way, the AI system provides a detailed and progressive analysis of the embryo’s development over a specific time period, starting only with the initial seven frames. The development of embryos differed significantly between those selected for transfer and those discarded.

The AI system effectively adjusted to these variations across different developmental cell stages of embryo morphology in both categories. To the best of our knowledge, the forecasting approach, predicting time-lapse video frames depicting the future of embryo development, is a novel approach. The main contribution of this study lies in providing the embryologists with a visual tool to observe the varying progression of embryo morphology in transfer and avoid videos over subsequent hours into the future. The visualization facilitates the identification of key biomarkers important for evaluating embryo quality at an early stage of development.

## Materials and methods

### Ethics statement

The study was conducted according to the guidelines of the Regional Committee for Medical and Health Research Ethics–South East Norway, and the General Data Protection Regulations. The embryo videos used in the study (for both training and evaluation) and the associated patient information were fully anonymized and collected after the approval by the Regional Committee for Medical and Health Research Ethics–South East Norway.

### Time lapse incubator

Time-lapse technology in human embryo incubation allows continuous monitoring of embryo development without affecting the conditions like temperature, pH, and gas concentration [11, 24]. A time-lapse incubator combines an IVF incubator with an integrated microscope and camera to capture pictures of embryo development every 5 to 20 minutes at several focal planes [25]. These images are then compiled into a video. The video is used for assessing embryo viability and annotating parameters, such as the start of the blastocyst. In this study, we used a time-lapse system called Embryoscope^TM^ by Vitrolife. The system is fitted with a plate referred to as an “embryoslide”. The embryoslide hosts several wells for culturing embryos individually. When a well enters the microscope’s field of view, an image is acquired by the inbuilt camera. The system used a camera with a LED light source (under 635 nm) passing through Hoffman’s contrast modulation optics. For each embryo, the system captured 8-bit images with a resolution of 250 *×* 250 pixels on different focal planes (usually between 3 or 5) at intervals of 7, 15, or 20 minutes. However, in the study we considered images from the central focal plane only. In this study, embryologists annotated each video for the embryo cell stage timings.

### Data

The dataset, used in this study, consisted of 365 time-lapse videos recorded at the fertility clinic Volvat Spiren in Oslo, Norway between 2013 and 2019. All 365 videos had a low rate of fragmentation, i.e., up to 15%. Videos with a rate of fragmentation above 15% were excluded from the dataset. Fragmentation refers to the cytoplasm content that remains outside the daughter cells during cell division, and the rate of fragmentation quantifies the percentage of such material within an embryo. We divided the 365 videos into two datasets. The first dataset comprised of 220 videos corresponding to day 2 of embryo development between 31 hpi to 43 hpi (12 hours) and is referred to as ‘cells stage study’. The second dataset had the remaining 145 videos representing day 4 of embryo development between 90 hpi to 113 hpi (23 hours), referred to as ‘blastocyst study’.

### Data preprocessing

We preprocessed the datasets in the cells stage study and the blastocyst study. As a first step of preprocessing, each video frame was resized from 250x250 to 128x128, normalized and additional image channels were stacked onto the frame. Fig 2 shows the data preprocessing workflow and stacking of the additional channels to the video frames. In the cells stage study, two additional channels represented the cell count and embryo development time, reported in hpi. For the blastocyst study, the additional channel represented only embryo development time. We omitted the cell count channel due to the difficulty in counting cells due to cell compaction. These additional channels information was derived from embryologist annotations accompanying the time-lapse videos. We normalized the channel information before stacking them with image channels.

**Fig 2.**
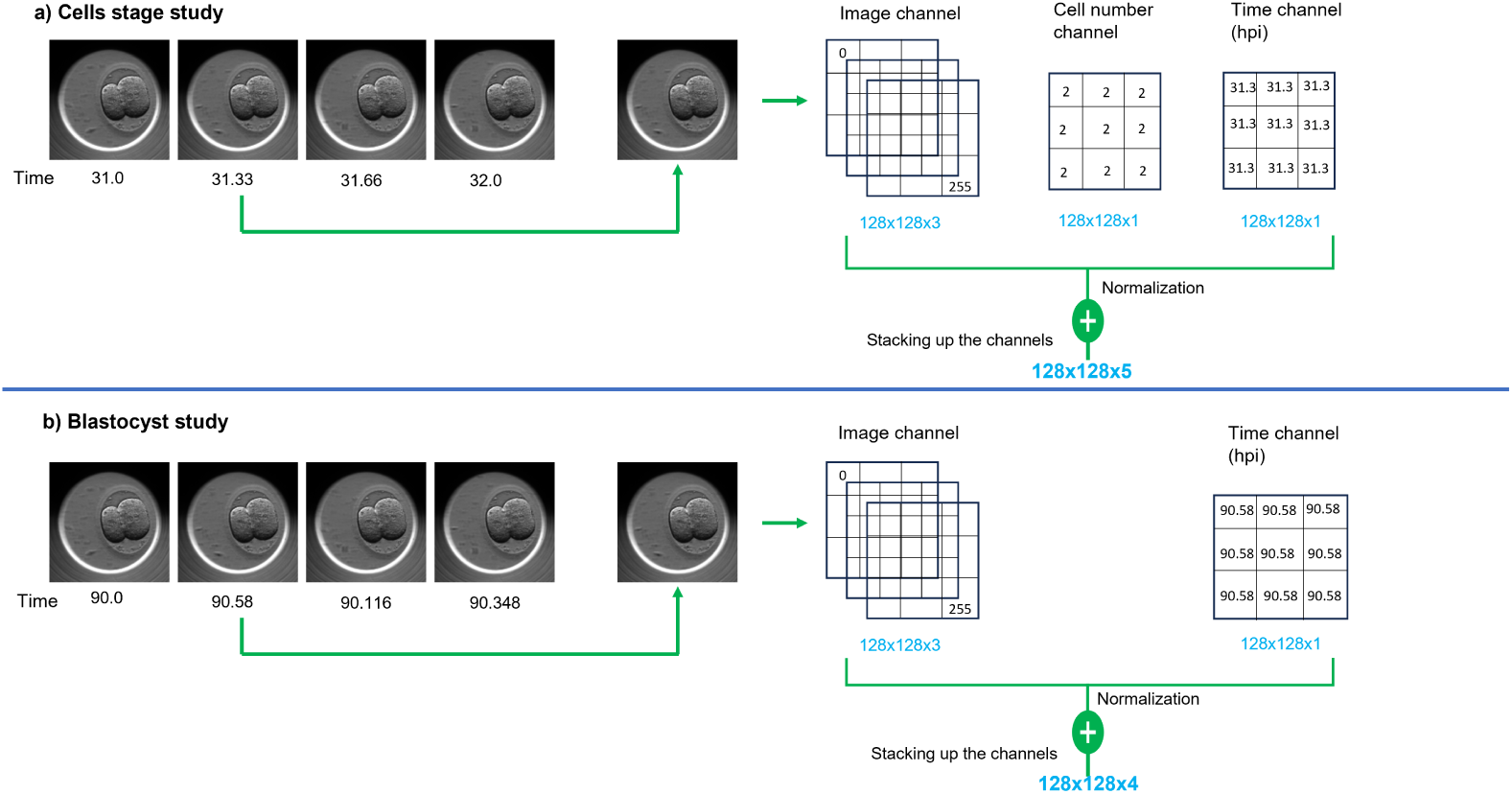
Preprocessing input frames of the embryo videos. A workflow diagram outlining the stacking of additional channels to the input frames of an embryo video. The channels represent development time (in hours post insemination) and cell count. Part a): Cells stage study: preprocessing a frame by stacking additional channels: cell count and time. Part b): Blastocyst study: preprocessing a frame by stacking only the time channel.

### Problem definition

Forecasting involves predicting the future frame sequence of a time-lapse embryo video. For sequence prediction, the initial step involves a single frame prediction task. The task aims to deduce the next video frame using the morphology from the previous frames. Let *X_t−_*_1_*_,F_* = [*x_t−F_ , . . . , x_t−_*_1_] represent a video sequence from time *t − F* to time *t −* 1, where the term *x_i_* ∈ ℝ*^C,H,W^ ∀i* corresponds to a video frame with channels *C*, height *H* and width *W* . The frame prediction model is responsible for predicting the next frame; *x_t_*. Let 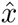*_t_* represent our prediction of *x_t_*. Let *Y_t,F_* = [*x_t−_*_(_*_F−_*_1)_*, . . . , x_t−_*_1_, 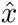*_t_*] denote the sequence shifted one time step and the prediction added. Let *G*_Θ_ define a parametric prediction model, providing a mapping function *G*_Θ_ : *X_t−_*_1_*_,F_ 1→ Y_t,F_* . The mapping function will be optimized by minimizing the difference between the observed sequence and the predicted sequence, where *ℓ* is the binary cross entropy loss function.

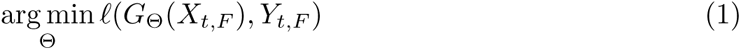

In this study, we built a video frame prediction model using Convolutional LSTM [26] layers. The model architecture is explained in Section Video frame prediction model.

### Convolutional LSTM

Recurrent neural network, explained in S1 Appendix, has introduced the concept of the hidden state for holding the network’s previous computations. The idea is refined by LSTM, explained in S2 Appendix, by introducing cell states *c_t_*and information processing using gates. Convolutional LSTM (ConvLSTM) [26] is an extension of LSTM used for spatio-temporal prediction. Similar to LSTM, ConvLSTM also passes information from the previous hidden state to the next frame of the video sequence, but the matrix multiplications defining state to state transitions are the local convolution operations defined on spatial axes. The mathematical equations and technical details are provided in S3 Appendix. Let *X_t_* be defined as a video frame, then ConvLSTM considers the pixels belonging to *X_t_* as the cells on a spatial grid. ConvLSTM captures changes in pixel intensity across this grid. Next, ConvLSTM determines the future of a grid cell based on the current inputs and the past LSTM states (hidden state, cell state) around its local neighborhood. Assuming *X_t_* is the third frame of a video sequence with the embryo 4-cell stage. The embryo was in the 2-cell stage in the first and 3-cell stage in the second frame. Now, ConvLSTM attempts to predict the next frame of the video (a 2D image matrix) based on the time-step progression of the image pixels’ convolutional feature vector in frame one, two and three. The inner structure of the ConvLSTM capturing the temporal dependencies in a video sequence is shown in S1 Fig. We used ConvLSTM to build the video frame prediction model because ConvLSTM has been widely applied for predicting the upcoming frames using previous frames [27–30].

### Video frame prediction model

We refer to our video prediction model as the ‘FramePredictor’ in the subsequent text. Fig 3 shows the architecture of the FramePredictor using a block diagram representation. The network has four ConvLSTM layers with ReLU activation. The layers have 32 filters, 64 filters, 128 filters and 128 filters with kernel sizes to be 7, 5, 3 and 1 respectively. The first three layers are followed by batch normalization with the ‘return sequence’ set as true. The last layer is followed by a dropout layer (rate=0.25) and has the ‘return sequence’ set to false. The ‘return sequence’ parameter in ConvLSTM layers determines the length of the output (number of video frames). If the parameter is true, the length of the input and the output sequence is the same. As the FramePredictor predicts a single frame, the ‘return sequence’ is set to false in the last ConvLSTM layer. The dropout layer’s output splits between two convolution layers, referred to as convHead_1_ and convHead_2_, as shown in Figure 3. Both convHead_1_ and convHead_2_ have a kernel size of 3, employs sigmoid activation functions for predictions.

**Fig 3.**
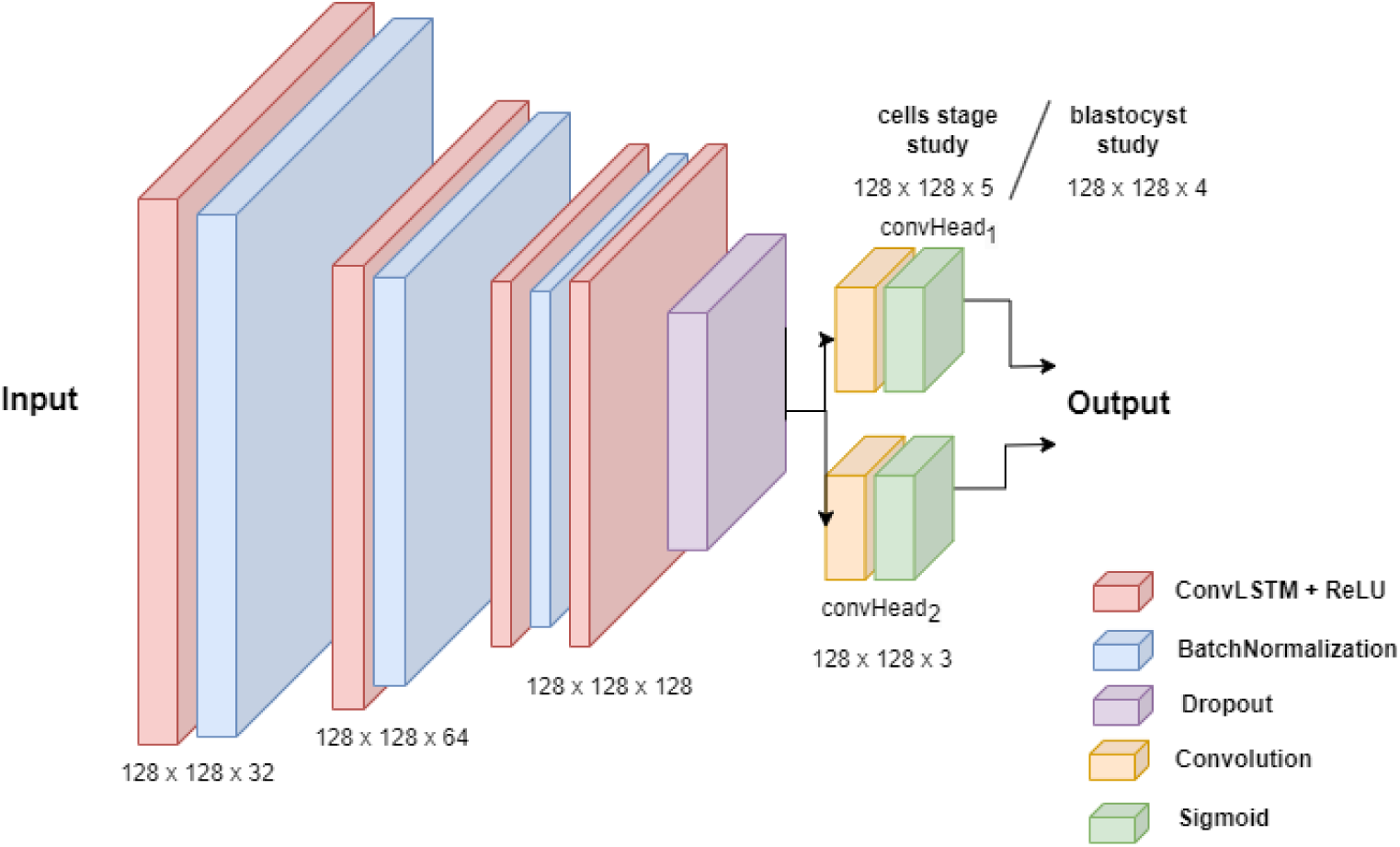
Architecture of the FramePredictor. A sequential model with ConvLSTM layers followed by a batch normalization layer or a dropout layer. The input consists of a video sequence and the output comprises two images. The first output’s image dimension varies depending upon the study type, but the dimension of the second output is constant across both studies.

For both the cells stage and blastocyst studies, we set *F* = 7 and the dimensions *W* = *H* = 128. However, the channel depth was set to *C* = 5 for the cells stage study and *C* = 4 for the blastocyst study. It resulted in convHead_1_ predicting a frame with dimensions of 128 x 128 x 4 for the blastocyst study and 128 x 128 x 5 for the cells stage study. Whereas convHead_2_ predicts a frame with dimensions of 128 x 128 x 3. We used the Keras library for constructing the layers of the FramePredictor and the architecture was inspired by the ConvLSTM model in Keras Code examples [31].

### Training of the FramePredictor

For training of the FramePredictor, the frames of a video are rearranged in a specific manner. Consider a video consisting of 12 frames. As FramePredictor uses an input sequence of seven frames length, the video is reorganized into several subsequences with overlapping frames. The first subsequence consists of frames *x*_1_ to *x*_7_, with the *x*_8_ frame as the ground truth. The second subsequence comprises frames *x*_2_ to *x*_8_, with the *x*_9_ frame as the ground truth. This pattern continues, with each subsequent sequence shifting by one frame until the last frame is used as the ground truth for training the FramePredictor. In this example, the last frame is *x*_12_.

The FramePredictor predicted the embryo morphology in time-lapse videos specifically for two periods: the cells stage study and the blastocyst study as explained in Section Data and Section Data preprocessing. The datasets comprises of two categories of videos: “transfer videos” of embryos transferred to females and “avoid videos” of embryos discarded by embryologists. We separately trained the FramePredictor on both transfer and avoid videos, resulting in four distinct models for prediction tasks: cells stage study with avoid videos (studyCA), cells stage study with transfer videos (studyCT), blastocyst study with avoid videos (studyBA), and blastocyst study with transfer videos (studyBT). Each study’s dataset was further divided with detailed descriptions provided in the following sections.

#### Dataset: cells stage study

Within this set, there were 110 transfer videos and 110 avoid videos. Among these 220 videos, 94 had a frame rate of 3 frames per hour, while 126 had a frame rate of 4 frames per hour. In all 220 videos, the first video frame always had the embryo in the 2-cell stage. We describe the exact cell stage data distribution within the dataset in S2 Fig.

We split the 110 avoid videos into two subsets: 100 for training the studyCA. The dataset is referred to as the trainCA set (<u>train</u> <u>c</u>ells <u>a</u>void). The remaining 10 referred to as the evalCA set (<u>eval</u>uation <u>c</u>ells <u>a</u>void) was used as an independent dataset for assessing studyCA. Similarly, the transfer videos were divided into two groups: 100 for training the studyCT, referred to as the trainCT set (<u>train</u> <u>c</u>ells transfer), and the remaining 10, referred as the evalCT set (<u>eval</u>uation <u>c</u>ells transfer) was used in the independent evaluation of the studyCT. Here, ‘independent dataset’ refers to videos that were not used in the model’s training or validation.

#### Dataset: blastocyst study

This dataset contained 71 transfer videos and 74 avoid videos, each with a frame rate of 8 frames per hour. The avoid videos had embryos beginning at the 9+ cell stage, whereas the transfer videos captured embryos from the start of cell compaction. Avoid videos started with embryos in the 9+ cell stage. Transfer videos started with embryos at the start of the compaction. We detail the distribution of embryo development stages across this dataset in S2 Fig.

Of the 74 avoid videos, 66 videos referred to as the trainBA set (<u>train</u> <u>b</u>lastocyst <u>a</u>void) were used in training the studyBA. The remaining 8 videos referred as the evalBA set (<u>eval</u>uation <u>b</u>lastocyst <u>a</u>void) was used as independent dataset for evaluating the studyBA. The 71 transfer videos were divided into two sets: 64 videos for training studyBT, referred to as the trainBT set (<u>train</u> <u>b</u>lastocyst transfer) and 7 videos for evaluating the studyBT, referred to as the evalBT set (<u>eval</u>uation <u>b</u>lastocyst transfer).

#### Training hyperparameters

The datasets trainCA, trainCT, trainBA and trainBT were divided into a ratio of 80:20 for training and validation testing of studyCA, studyCT, studyBA and studyBT respectively.

For the cells stage study, we processed input sequences of size 7x128x128x5, producing two output images with the sizes 128x128x3 and 128x128x5 or 128x128x4. The training involved 50 epochs with a batch size of 2, utilizing the Adam optimizer with a learning rate 10*^−^*^4^ and binary cross entropy as the loss function. During training, we saved the model based on the highest validation accuracy, specifically for predicting the 128x128x3 images. For the blastocyst study, the input dimensions were 7x128x128x4 and the output consisted of two images with dimensions of 128x128x4 and 128x128x3. The training involved 35 epochs and had the other parameters to be the same as described in the cells stage study.

### Embryo cropper

The predicted video frames have an embryoslide well visible in the background. However, because our primary focus is on the embryos (foreground), we removed the embryoslide wells from both the original and predicted video frames during the evaluation of the FramePredictor. Since we made many predictions, we automated the cropping by training a U-Net model, referred to as the ‘embryo cropper’ for generating the masks to separate embryos from the background. Fig 4 shows an example of a video frame after post-processing with the embryo cropper.

**Fig 4.**
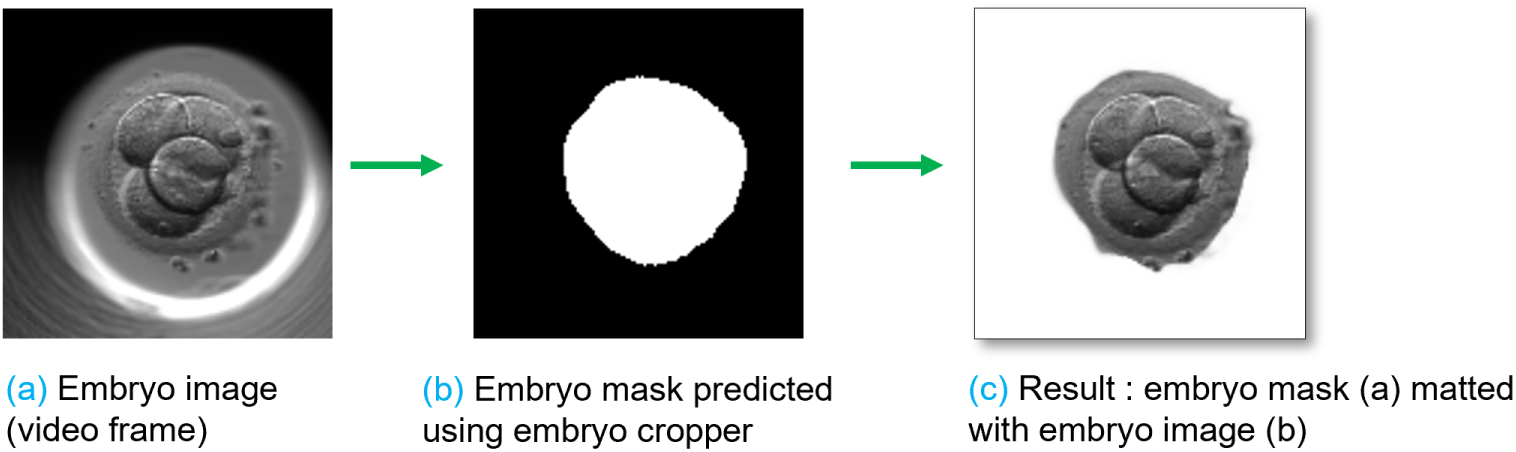
Post-processing with the embryo cropper. Part a): A frame extracted from a time-lapse embryo video. Part b): Embryo cropper locates the embryo region and predicts the segmentation mask. Part c): Embryo cropper mattes the mask (b) to the image (a) for extracting the embryo region.

#### Dataset: embryo cropper

Based on empirical testing, we used 1994 embryo images (video frames), extracted at random time intervals from the 365 videos (Section Data). This dataset was referred to as the ‘trainCrop’ (<u>train</u> embryo cropper) dataset. Within the dataset, the distribution of embryo cell stages was as follows: 250 images each for the 2-cell, 3-cell, 4-cell, 9+ cell, start of compaction, morula, start of blastocyst stages, and 244 images for the full blastocyst stage. We used LabelBox [32] to annotate the dataset for obtaining the segmentation masks used for training the embryo cropper.

#### Training: embryo cropper

We used the standard vanilla U-Net architecture and trained it on the trainCrop dataset. The input image size was set to 128x128x3, and we applied both horizontal and vertical flipping as data augmentation techniques. The dataset was divided into training and validation set at the ratio of 80:20. The batch size used was 32, the number of epochs was 60. We used Adam optimizer with a learning rate of 10*^−^*^3^. The loss function was sparse categorical cross entropy. The trained embryo cropper generates a mask to crop out the embryo region from a video frame. That same mask is matted with the frame (ground truth) and the predicted frame to extract the embryo region. We used the vision transformer VitMatte [33, 34] for the matting process.

### Forecasting embryo development

For a video sequence as an input, the FramePredictor is capable of only predicting the next immediate frame following the last frame. However, the aim of this study is to forecast embryo development over a couple of hours into the future, not just single frame prediction. Hence, we devise two forecasting strategies based on the predictions from the FramePredictor. The strategies are referred to as ‘Forecasting till the end’ and ‘forecasting the next 7 frames’. Fig 5 shows a schematic representation of the workflow for both the forecasting strategies. The frame sequences from a video are reorganized into subsequences, following a similar rearrangement scheme as explained in Section Training of the FramePredictor. The rearrangement is conducted irrespective of the chosen forecasting strategy.

**Fig 5.**
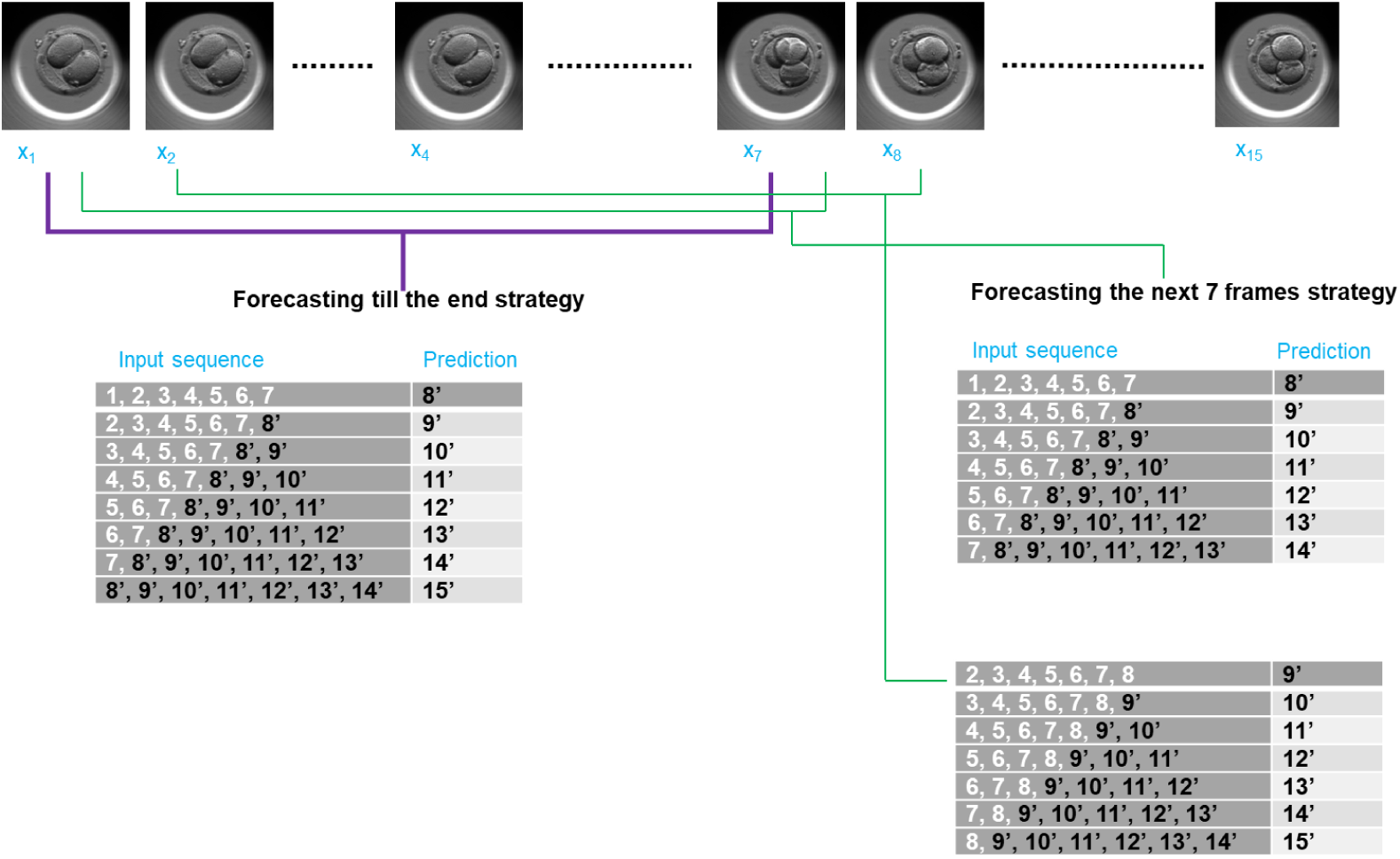
Forecasting embryo development. Considering an embryo video with 15 frames. Each frame is referenced with its location in the video: Frame *x*_1_ becomes 1 and predicted frame *x*^_8_ becomes 8*^′^*. ‘Forecasting till the end’ uses the first seven frames to forecast the remaining frames (8*^′^* to 15*^′^*). The strategy appends 8*^′^* to the input for forecasting 9*^′^* and continues so forth. ’Forecasting the next 7 frames’ uses the first seven frames but forecasts only the next seven frames (8*^′^* to 14*^′^*). Later, when frame 8 is received from the embryo video, the strategy uses frames 2 to 8 to update the forecasts of frames 9*^′^*to 14*^′^* and in addition makes a forecast of frame 15*^′^*.

In the Forecasting till the end strategy, we modify the input sequences by including the predicted frames within the sequence itself. This modified input is then used by the FramePredictor to forecast subsequent video frames. For example, for an input sequence with frames: *x*_1_, *x*_2_, *x*_3_, . . . , *x*_6_, *x*_7_, the FramePredictor predicts the next frame *x*^_8_. The forecasting scheme will transform the input sequence into *x*_2_, *x*_3_, *x*_4_, . . . , *x*_7_, 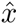_8_ for predicting 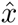_9_. The process continues with the predictions (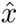_10_, 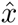_11_, and so on) until we reach the end of both studies.

Forecasting the next 7 frames strategy involves predicting the subsequent seven frames only. The approach is based on the idea that with each new frame obtained from the incubator for a time-lapse video, the system updates its forecast for the next seven frames. Given an initial sequence of frames (*x*_1_, *x*_2_, *x*_3_, . . . , *x*_6_, *x*_7_), the strategy will forecast the upcoming seven frames (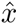_8_, 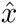_9_, 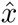_10_, . . . , 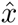_13_, 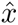_14_). After predicting 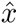_14_, the prediction is paused until frame *x*_8_ is acquired. The pause period depends on the video’s frame rate. The input sequence changes to *x*_2_, *x*_3_, *x*_4_, . . . , *x*_7_, *x*_8_. Now, using the modified input sequence, the forecasting process continues by updating the forecasts of 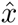_8_, . . . , 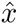_14_ and in addition making the forecast 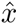_15_. The scheme continues to forecast until the end of the studies is reached.

### Performance metrics

To evaluate the proposed system, we use state-of-the-art video quality evaluation metrics. The metrics included Peak Signal to Noise Ratio (PSNR), structural similarity index measure (SSIM) [35] and Fŕechet Video Distance (FVD) [36]. We use SSIM and PSNR because of their effectiveness in measuring the deviation of the predicted sequence from the baseline ground truth sequence given a specific scenario [36]. In our case, the scenario is related to differences in embryo morphology. Essentially, these metrics penalize predictions that deviate from the ground truth [37]. Thus, we could effectively quantify the degree of divergence in the predicted embryo morphology. We used scikit-image [38] for calculating SSIM and PSNR.

The FVD score extends the underlying concept of Frechet Inception Distance [39].

The FVD metrics take into account the temporal consistency between two videos (ground truth and predicted). Besides measuring the video quality [36] and a lower FVD score is indicative of higher video quality. By calculating the FVD scores we investigated the maximum duration for which the dynamics of the predicted sequence align with the ground truth.

## Results

We evaluated the performance of the FramePredictor. Later, we evaluated the performance of the FramePredictor with the two forecasting strategies. Below, we present the evaluation results for both studies: the cells stage study and the blastocyst study.

### Evaluation of FramePredictor

We evaluated the performance on studyCA, studyCT, studyBA, and studyBT using both validation testing sets (from trainCA, trainCT, trainBA and trainBT) and independent sets (evalCA, evalCT, evalBA and evalBT). We computed PSNR and SSIM metrics on a per-frame basis and averaged them, but the FVD score was computed for sequences of frames. We based metric calculations on the predictions with a dimension of 128x128x3. For evaluation, we reorganized video sequences into sub-sequences using the sliding window technique outlined in Section Training of the FramePredictor.

The per-frame evaluation results are reported in Table 1, which includes outcomes before and after applying the embryo cropper post-processing. We observed consistent PSNR and SSIM values across both the validation and independent sets for the transfer and avoid videos, except in the avoid category’s blastocyst study. There was a drop in performance numbers from validation to independent set, indicating the FramePredictor’s limited generalization for studyBA. For transfer videos, both PSNR and SSIM reported slightly higher value in comparison to avoid videos. The metric values further increased after performing the post-processing with embryo cropper, suggesting that FramePredictor focuses on relevant embryo features to predict morphological changes. The highest performance was observed in the transfer videos of the independent set following embryo cropper post-processing.

**Table 1.**
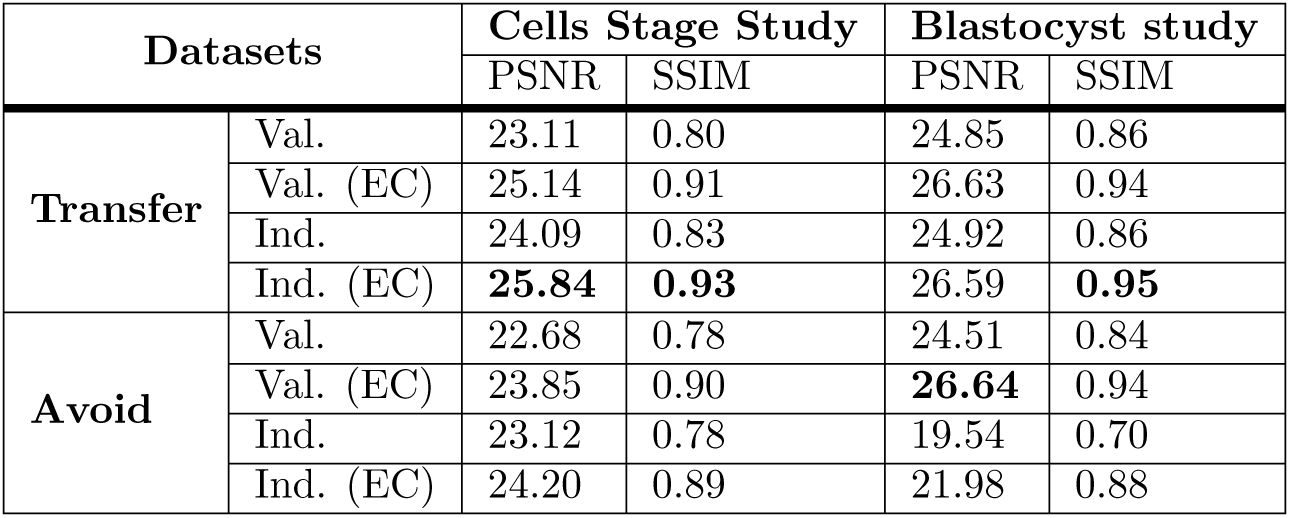
Per-frame evaluation of the FramePredictor’s predictions for cells stage and blastocyst studies. PSNR and SSIM metrics, calculated for both the validation (Val.) and independent (Ind.) sets across transfer and avoid videos, are presented with and without embryo cropper (EC) post-processing. The best results are shown in bold.

| Datasets |  | Cells Stage Study |  | Blastocyst study |  |
| --- | --- | --- | --- | --- | --- |
|  |  | PSNR | SSIM | PSNR | SSIM |
| **Transfer** | Val. | 23.11 | 0.80 | 24.85 | 0.86 |
|  | Val. (EC) | 25.14 | 0.91 | 26.63 | 0.94 |
|  | Ind. | 24.09 | 0.83 | 24.92 | 0.86 |
|  | Ind. (EC) | **25.84** | **0.93** | 26.59 | **0.95** |
| **Avoid** | Val. | 22.68 | 0.78 | 24.51 | 0.84 |
|  | Val. (EC) | 23.85 | 0.90 | **26.64** | 0.94 |
|  | Ind. | 23.12 | 0.78 | 19.54 | 0.70 |
|  | Ind. (EC) | 24.20 | 0.89 | 21.98 | 0.88 |

We assessed development using the FVD score at three different time intervals for the cells stage study: 2-3 hours with 10 frames, 4-5 hours with 14 frames, and 7-10 hours with 28 frames. In the blastocyst study, we analyzed videos trimmed to lengths of 60 frames (8 hours), 120 frames (15 hours), and 180 frames (22 hours). The duration was calculated based on the videos’ frame rate. We calculated FVD score on the independent sets only because the validation set had frame sequences covering partial portions from different videos and cannot be evaluated for representing a complete video context. We calculated the FVD score only on independent sets because the validation set consists of frame sequences that represent only partial segments from different videos. These sequences do not provide a full video context, therefore unsuitable for evaluation.

The FVD score at different video lengths is reported in Table 2. The table also includes the results after the predictions were post-processed with the embryo cropper.

**Table 2.** Evaluating the FramePredictor’s predictions of video context for cells stage and blastocyst studies. FVD scores, calculated the independent (Ind.) set across transfer and avoid videos, are presented with and without embryo cropper (EC) post-processing. The videos were trimmed at different length with F number of frames. The best results are shown in bold.

| Datasets |  | FVD score |  |  |  |  |  |
| --- | --- | --- | --- | --- | --- | --- | --- |
|  |  | Cells stage study |  |  | Blastocyst study |  |  |
|  |  | F=10 | F=14 | F=28 | F=60 | F=120 | F=180 |
| **Transfer** | Ind. | 2473.81 | 1721.33 | **904.39** | 779.18 | 380.18 | **236.33** |
|  | Ind. (EC) | 722.11 | 526.43 | **355.62** | 343.99 | 247.97 | **173.33** |
| **Avoid** | Ind. | 1869.71 | **1441.56** | 1467.33 | 740.80 | 391.17 | **223.98** |
|  | Ind. (EC) | 633.91 | **536.44** | 582.38 | 376.42 | 232.03 | **165.13** |

We observed that FVD scores decreased as the video sequences became longer, indicating an improved prediction quality with a higher coverage of embryo development. Thus, the FramePredictor accurately predicts the embryo’s development over longer periods. The highest performance was observed in the 180-frame sequences of the blastocyst study. For the cells stage study, the best FVD score was in the28-frame sequences for transfer videos and in the 14-frame sequences for avoid videos. Post-processing with the embryo cropper led to a significant decrease in FVD scores. This suggested that the background artifacts might have contributed to the context mismatch between predictions and ground truth.

Fig 6 shows a video sequence predicted by FramePredictor in the transfer category for both the cells stage and blastocyst studies. Upon assessing the image quality of the predictions, we observed that the visual quality of the predicted frames was lower in comparison to the ground truth frame, across both the studies. By the term ”quality,” we referred to embryo images with clear cell membranes, lack of noise or artifacts. For the cells stage study, predicted frames sometimes had the presence of artifacts such as gray-scale color distortions in the background or around the zona pellucida. The zona pellucida is a circular ring surrounding the embryo region. For the blastocyst study, the initial predictions had the presence of gray-scale color distortions in the background.

**Fig 6.**
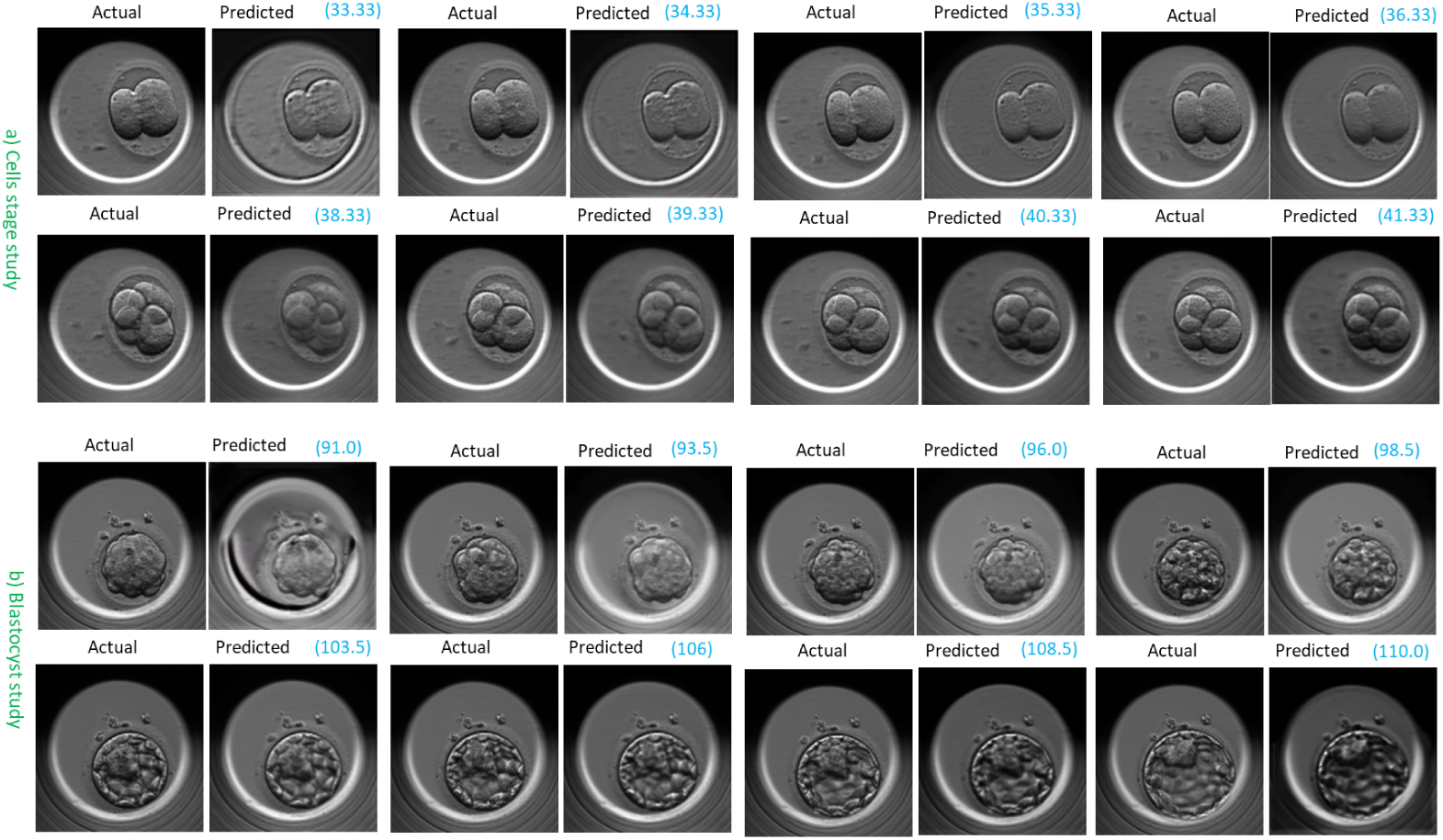
FramePredictor’s predicted embryo development at various time points. a) A transfer video from the independent set (evalCT) in the cells stage study b) A transfer video from the independent set (evalBT) in the blastocyst stage study. The actual (ground truth) frames are on the left, while the predicted frames are on the right, with time points highlighted in blue and reported in hours post insemination.

The blurred cell membranes was another issue. For the cells stage study, the cell membranes were always visible, but, the blurriness and artifacts reduced the overall visual appeal of the predictions. These issues with the image quality were more pronounced for the predictions in the avoid category. We present a few more examples of the FramePredictor’s predictions for cells stage study in S3 Fig and for the blastocyst study in S4 Fig.

Regardless of the video category, the image quality at the beginning was much lower when compared to other segments of the video. The quality improved significantly towards the end segment of the video.

#### Assessment by the embryologists

The predicted frames for the cells stage study and the blastocyst study were assessed by four embryologists. According to their assessment of the predictions, transfer videos provided enhanced visibility of embryo morphology over avoid videos. Few predicted frames in the transfer category had structures similar to cell nuclei. However, validating these predictions for clinical use is undermined by the presence of gray-scale distortions around the zona pellucida and the difficulty in distinguishing individual cells due to blurred cell membranes. But, the predictions were effective in signaling the beginning of the blastocyst stage.

### Evaluation of the forecasting strategies

We re-evaluated the FramePredictor on independent sets using the exact settings as described in Section Evaluation of FramePredictor, but the predictions were recursively utilized by the forecasting strategies to further predict the embryo’s development. Below, we present evaluation results for the FramePredictor with both forecasting strategies.

#### Forecasting till the end

The evaluation of the ‘Forecasting till the end’ strategy is reported in Table 3, which includes outcomes before and after applying the embryo cropper post-processing. For the cells stage study, PSNR and SSIM metrics were higher, whereas the blastocyst study reported lower FVD scores. The strategy performed better for transfer videos than for avoid videos. Post-processing the predictions with the embryo cropper resulted in improving the metrics values across all datasets.

**Table 3.** Evaluating forecasting strategies on independent datasets of cells stage study. Table reports PSNR, SSIM and FVD score (FD) for ‘Forecasting till the end’ before and after the application of embryo cropper (EC) post-processing on independent (Ind.) sets. The best results are shown in bold.

| Datasets |  | Cells stage study |  |  | Blastocyst study |  |  |
| --- | --- | --- | --- | --- | --- | --- | --- |
|  |  | PSNR | SSIM | FVD | PSNR | SSIM | FVD |
| Transfer | Ind. | 14.51 | 0.40 | 3728.40 | 11.19 | 0.14 | 2882.38 |
|  | Ind. (EC) | **18.02** | 0.74 | **1159.90** | **13.75** | 0.64 | 1822.02 |
| Avoid | Ind. | 11.77 | 0.20 | 4953.18 | 10.70 | 0.22 | 3766.75 |
|  | Ind. (EC) | 16.23 | **0.76** | 1983.22 | 13.24 | **0.69** | **1743.93** |

#### Forecasting the next 7 frames

The evaluation results of the ‘Forecasting the next 7 frames’ strategy is reported in Table 4. The table also includes the results after the predictions were post-processed with the embryo cropper. Metric analysis revealed that the ‘Forecasting the next 7 frames’ strategy outperformed the ‘Forecasting till the end’ approach. In comparison to the latter, the former had higher PSNR and SSIM, and a lower FVD score. This is as expected, since forecasting to the end is a more challenging problem than forecasting just the next seven frames.

**Table 4.** Evaluating ‘Forecasting the next 7 frames’ strategy on independent datasets of cells stage study. Table reports PSNR, SSIM and FVD scores (FD) for ‘Forecasting the next 7 frames’ before and after the application of embryo cropper (EC) post-processing on independent (Ind.) sets. The best results are shown in bold.

| Datasets |  | Cells stage study |  |  | Blastocyst study |  |  |
| --- | --- | --- | --- | --- | --- | --- | --- |
|  |  | PSNR | SSIM | FVD | PSNR | SSIM | FVD |
| Transfer | Ind. | 12.90 | 0.53 | 4449.18 | 15.81 | 0.51 | 833.04 |
|  | Ind. (EC) | **19.40** | **0.82** | **966.27** | **19.40** | **0.82** | **433.79** |
| Avoid | Ind. | 11.15 | 0.43 | 4843.87 | 17.70 | 0.57 | 1282.08 |
|  | Ind. (EC) | **17.18** | **0.77** | **1186.65** | **19.10** | **0.82** | **791.27** |

The performance of the ‘Forecasting the next 7 frames’ strategy improved significantly after applying embryo cropper post-processing, with the highest metric scores recorded post-processing in both the cells stage and blastocyst studies. This enhancement was consistent across both transfer and avoid videos. Additionally, the forecasting strategy performed better for transfer videos.

Fig 7 shows the ‘Forecasting the next 7 frames’ strategy’s predicted sequence for the transfer category in the cells stage and blastocyst study. Since the forecasting approach used predictions from FramePredictor as an input, there was noticeable grayscale level distortions in the forecast. Similarly, the blurriness associated with cell membranes was also more pronounced. Furthermore, we compared the forecasted embryo development to the embryologists’ annotations for the embryo development in the ground truth videos. The forecasting strategy typically had an average time delay of 1 to 1.5 hours in predicting transitions between cell stages.

**Fig 7.**
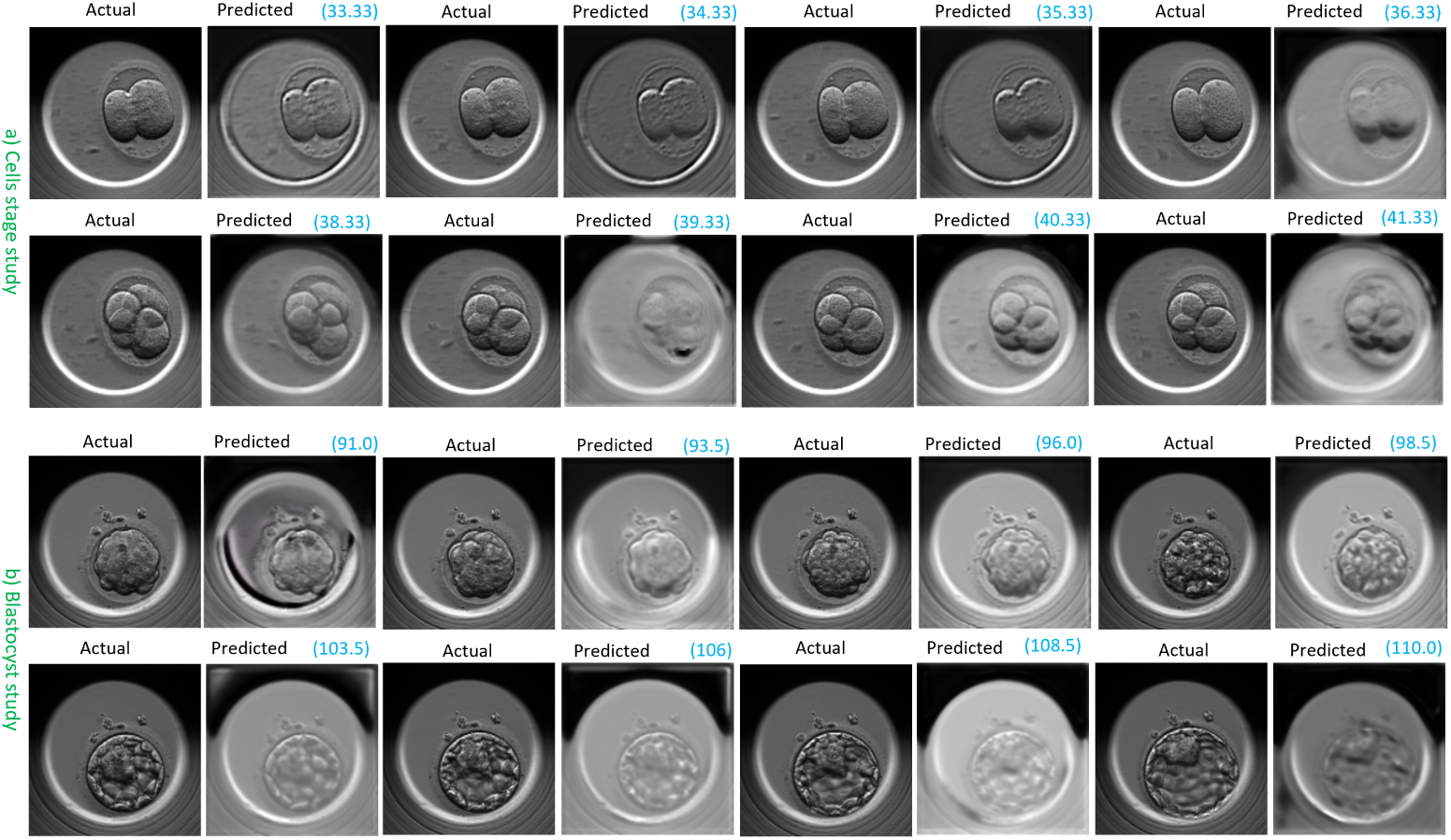
Forecasted embryo development at multiple time points with the ‘Forecasting the Next 7 Frames’ strategy. a) A transfer video from the independent set (evalCT) in the cells stage study b) A transfer video from the independent set (evalBT) in the blastocyst stage study. The actual (ground truth) frames are on the left, while the predicted frames are on the right, with time points highlighted in blue and reported in hours post insemination.

#### Assessment by the embryologists

The forecast from the ‘Forecasting the next 7 frames’ strategy was also evaluated by same embryologists. Noise such as artifacts, distortions, and blurred cell membranes made it challenging for them to confidently determine the beginning of cell stages in forecasts, although transfer videos were more clearer than avoid videos. Embryologists could observe changes within the embryo in forecasted sequences, but for clinical validation, a noise-free prediction is necessary. In the cells stage study, forecasts between 36 to 38 hpi and between 40 to 41 hpi were exceptionally clear. The start of the blastocyst could always be accurately identified in the blastocyst study, but the transition to a full blastocyst with an inner cell mass was not as clear. For avoid videos, noise and blurriness prematurely suggested cell or blastocyst degeneration and sometimes cells were mistaken for large fragments, but correct differentiation was possible upon reviewing longer in the forecast.

## Discussion

The AI system demonstrated a strong understanding of embryo morphology dynamics. The system’s FramePredictor predicted video frames that accurately captured the developmental context of embryos in both the cells stage and blastocyst studies, across ’transfer’ and ’avoid’ video categories. The FramePredictor was particularly more effective in the transfer category. Furthermore, post-processing the predicted frames with the embryo cropper to eliminate background led to significant improvements in video quality metrics (PSNR, SSIM, and FVD scores). This indicates that the predicted outcome focused on the relevant regions of the embryo rather than merely replicating the background across frame sequences.

The quality of the predicted frames did not meet the standard of the ground truth. At times, the predicted morphology was blurry and had grayscale color distortions. Consequently, this lowered the chance of clinically validating the predictions. These distortions were primarily due to adding extra channels to the input frame sequences. The additional channels provided time period and embryo cell count information enabling the FramePredictor to better understand embryo morphology changes. However, this additional information affected the image feature distribution, leading to image distortions. A potential future research direction could involve revising the method of integrating this information. An alternative strategy could be to embed these channels in the latent space and evaluate the impact on image quality.

The AI system implemented two forecasting strategies for predicting embryo morphology changes: ‘Forecasting till the end’ and ‘Forecasting the next 7 frames’, across 12 hours in the cells stage study and 23 hours in the blastocyst study. The strategy ‘Forecasting the next 7 frames’ was more effective than the ‘Forecasting till the end’ strategy for both transfer and avoid videos. In the cells stage study, the ‘Forecasting the next 7 frames’ strategy provided valuable insights into the future of the embryo’s development on day 2. It included information such as whether an embryo divides by the end of the study, the synchronization of cell division, and the timing between adjacent cell stages. For the blastocyst study, the ‘Forecasting the next 7 frames’ strategy successfully determined whether the embryo will start to blastulate or not, which is crucial for assessing the embryo’s viability on day 4. However, forecasts were blurred, a limitation that could be improved with further training of the FramePredictor.

## Conclusion

The AI system accurately forecasted embryo morphology dynamics on day 2 (cells stage study) and day 4 (blastocyst study) of development. Using a sequence of seven video frames, it predicted the next seven frames, projecting two hours into the future of embryo development. The system effectively forecasted the progression of embryo morphology in both ’transfer’ and ’avoid’ categories. By separating the embryo from the background in the forecast and evaluating the predicted morphology, we demonstrated the AI system’s focus on crucial embryo features. In the ‘transfer’ category videos, the AI system accurately predicted the start of the blastocyst stage, with forecasts showing clearer cell membranes, fewer image gray-scale distortions and artifacts than in ’avoid’ videos. Despite initial issues with blurriness and distortions, the morphological clarity in forecasts improved significantly as the sequence progressed.

Upon evaluating the forecasts, embryologists concluded that changes in embryo morphology were detectable through the forecast sequence, but, image quality required enhancement for clinical validation. Therefore, the proposed AI system demonstrated a potential in assisting embryologists by providing insights into the future dynamics of embryo development.

The current study focused solely on the capabilities of a discriminative AI model in forecasting embryo development. As a next step in research, we plan to explore the use of generative AI models [40] for forecasting the embryo development.

## Author Contributions

### Conceptualization

Akriti Sharma, Alexandru Dorobantiu, Hugo L. Hammer

### Data curation

Akriti Sharma, Erwan Delbarre

### Formal analysis

Akriti Sharma

### Funding acquisition

Hugo L. Hammer, Mette H. Stensen, Erwan Delbarre, Michael A. Riegler

### Investigation

Akriti Sharma

### Methodology

Akriti Sharma, Alexandru Dorobantiu, Hugo L. Hammer

### Project administration

Akriti Sharma, Hugo L. Hammer

### Software

Akriti Sharma, Saquib Ali

### Supervision

Hugo L. Hammer, Mette H. Stensen, Erwan Delbarre, Michael A. Riegler

### Validation

Akriti Sharma, Mario Iliceto, Mette H. Stensen

### Visualization

Akriti Sharma

### Writing – original draft

Akriti Sharma

### Writing – review & editing

Akriti Sharma, Hugo L. Hammer, Mette H. Stensen, Mario Iliceto, Michael A. Riegler, Alexandru Dorobantiu

## Supporting information

**S1 Fig. Inner structure of Convolutional LSTM (ConvLSTM)** ConvLSTM processes an input matrix (*X_t_*) by modeling spatial distribution with temporal dependencies. ConvLSTM achieves this task through element-wise multiplication (hadamard product) of the feature vector of *X_t_* with the LSTM states at time t. *X_t_* corresponds to a video frame at t.

**S2 Fig. Embryo cell stages distribution in the datasets.** The videos used for training and evaluating the AI system contained distributions of embryo development stages as: Part a): Cell stages in the transfer videos for the cells stage study. Part b): Cell stages in the avoid videos for the cells stage study. Part c): Embryo development stages in the transfer videos for the blastocyst study. Part d): Embryo development stages in the avoid videos for the blastocyst study.

**S3 Fig. Cells stage study: FramePredictor’s predicted embryo development at various time points.** a) A transfer video from the independent set (evalCT). b) and c) Avoid videos from the independent set (evalCA). The actual (ground truth) frames are on the left, while the predicted frames are on the right, with time points highlighted in blue and reported in hours post insemination.

**S4 Fig. Blastocyst study: FramePredictor’s predicted embryo development at various time points.** a) A transfer video from the independent set (evalBT). b) and c) Avoid videos from the independent set (evalBA). The actual (ground truth) frames are on the left, while the predicted frames are on the right, with time points highlighted in blue and reported in hours post insemination.

**S5 Fig. Cells stage study: forecasted embryo development at multiple time points with the ‘Forecasting the Next 7 Frames’ strategy.** a) A transfer video from the independent set (evalCT). b) and c) Avoid videos from the independent set (evalCA). The actual (ground truth) frames are on the left, while the predicted frames are on the right, with time points highlighted in blue and reported in hours post insemination.

**S6 Fig. Blastocyst study: forecasted embryo development at multiple time points with the ‘Forecasting the Next 7 Frames’ strategy.** a) A transfer video from the independent set (evalBT). b) and c) Avoid videos from the independent set (evalBA). The actual (ground truth) frames are on the left, while the predicted frames are on the right, with time points highlighted in blue and reported in hours post insemination.

**S1 Appendix. Recurrent neural network: information processing, mathematical definitions and formulas**

**S2 Appendix. Long short-term memory: Information processing, mathematical definitions and formulas**

**S3 Appendix. Convolutional long short-term memory: Information processing, mathematical definitions and formulas**

